# Comparing apples to oranges from photoreceptors to inferior temporal cortex through neural manifolds

**DOI:** 10.64898/2026.10.30.763101

**Authors:** Yi Xu, Akihito Maruya, Bevil R. Conway, Qasim Zaidi

## Abstract

In the neural processing of color, cone photoreceptor outputs are combined over successive stages to form inferior temporal cortex (IT) cells with narrower color tuning and more diverse peak preferences. To understand the benefits of these transformations, we explored the capability of photoreceptor, retinal ganglion cell and IT population responses to distinguish between pairs of natural fruits and vegetables, based solely on their color distributions extracted from hyperspectral images. Simulations with realistic Poisson response variability show that linear separations are possible for many more pairs by IT neurons. Shapes of low-dimensional manifolds extracted from neural population responses explain the improvement.

## 1. INTRODUCTION

Human and macaque color vision begins with light absorption in three cone photopigments whose spectral sensitivities peak near 420, 530 and 560 nm and have broad, overlapping spectral tuning. Thomas Young’s original idea was given an empirical foundation by James Clerk Maxwell showing that a color-matching operation embedded all possible visible lights in a three-dimensional linear space formed by the three fixed primary spectral lights as axes, and that photoreceptor spectral sensitivities could be derived as linear transformations of the space defined by the triple of primary lights [1,2]. Maxwell also realized that filling the space required mechanisms that compared the outputs of the photoreceptors. In other words, the axes that represent the responses of the photoreceptors form just a 3-D “skeleton”, and to form a 3-D “space” requires cells that signal relative responses along more than one axis [3].

Much of color theory is implicitly about how points in the 3-D space representing combinations of cone responses are transformed to colors. Classically, opponent color theories from Hering, Schrödinger etc. purported to do this, but the lack of empirical evidence for a physiological or perceptual underpinning [4,5] has reduced opponent unique hues to the role of volitionally used compass-points in judging similarity between colors [6]. After the cone signals are processed through multiple retinal, thalamic and cortical stages, discrete loci in inferior temporal (IT) cortex form the last purely sensory areas for color. Color-sensitive neurons have been described in many extra-striate visual areas, notably within anterior (AIT) [7,8] and posterior (PIT) inferior temporal cortex [9]. Such neurons often respond selectively to small and circumscribed portions of the 3-D space, showing narrower tuning than photoreceptors and peak response at many different locations in the space. Using fMRI in macaques, Conway and Tsao [9] found color biased regions called “globs” in PIT, and using fMRI guided single-unit recording of 228 cells using 45 isoluminant colors equally spaced on a triangle in CIELUV space, they found cells with peak responses to 42 out of the 45 stimulus colors, each with significant responses to only a few neighboring colors, showing much narrower color tuning than receptors [10] or Retinal Ganglion Cells [11]. The progression from three broad and overlapping receptor sensitivities to a much larger diversity of distinct narrowly tuned curves raises questions about costs and benefits of the diversification and narrowing of color responses that have not been sufficiently addressed.

The limited overlap between IT tuning curves opens the possibility of IT population responses being the neural locus of color decoding [12]. Zaidi *et al*. [13] and Zaidi and Conway [3] used simulations to show that the diversity of preferred colors and the narrow tuning combined to enable identification of single colors with a simple winner-take-all decoding, even in the presence of Poisson response variability. This is consistent with a human observer reporting a blue-purple color near the center of gaze when stimulated by an electrode in a region of V4a that gave the largest field potentials to blue-purple stimulus colors [14]. Natural objects and surfaces have generic color variations, and color variations are an important cue in identifying objects and materials and their state [15–19]. Here we present computational simulations to test whether the benefits of diversity and narrowness extend to an increase in accuracy with which natural objects can be classified based on their color distributions. To answer this question, we compared the separation of color manifolds at photoreceptor (LMS), retinal ganglion cell (RGC) and PIT stages of the visual pathway. Using a hyperspectral database of 42 fruits and vegetables [20], which provides the wavelength spectrum for every pixel, we computed LMS, RGC and IT cell responses to each pixel of the fruits and vegetables. These formed population response distributions in high-dimensional neural ambient spaces, which we examined for separation and lower dimension manifolds.

In neuroscience and machine learning, manifolds are mathematical descriptions of coordinated neuronal population activity [21]. The term is borrowed from mathematics, where a manifold is a topological space of which every local neighborhood maps continuously to a Euclidean surface. If a manifold is smooth, then calculus can be used, and if it is Riemannian, a metric tensor can be used to define angles and distances. In neuroscience and deep neural networks (DNN), a manifold is a lower dimensional geometric object that embeds in a high dimensional ambient space representing activity of the neural population and is usually extracted by dimension-reduction methods that may need to be non-linear. Manifolds are useful conceptual devices in the cases where their geometry is preserved across individuals in the region performing the behavior. Manifolds have been used more widely in decoding neural control of motor behavior. For example, circular head rotation in mice yields a ring attractor manifold in the thalamus [22], and grid cell activity forms a donut-shaped toroidal manifold in the medial entorhinal cortex. The trajectory of exploration lies continuously on the surface of the manifold [23], and after learning, the shape of the manifold in cortical area CA1 matches the shape of the learned environment [24]. Manifolds have also been useful in understanding many computational principles used by the brain and DNNs. For example, population responses in motor cortex during reach preparation and movement, form two orthogonal subspaces dedicated to preparatory and movement computations, so the same neural population acts, at different times, as two separate circuits [25]. In a different vein, object categorization properties of DNNs have been shown to arise from repeated applications of the rectification operator ReLU which modifies the topology of the manifolds to reduce their Betti numbers, so that manifolds that are intertwined at early stages become hole free and thus separated at later stages [26]. Since rectification is ubiquitous in the visual brain, this result has direct implications for neural models of visual classification. Rectification would be integral to every model that explains the narrowing of color-tuned IT cells [27].

Since color vision begins with a 3-D ambient neural space with broad tuning curves and expands to a high-dimensional neural space with narrow tuning curves, some mathematical connections between tuning curves and manifolds that have been established under restricted conditions [28] may prove useful: Continuous tuning curves define a continuous mapping from stimulus properties to neural population response. The set of all population responses generated by the stimulus domain forms the neural response manifold. By definition, the mapping from stimulus property to the neural response is onto this manifold (surjective), since each point on the manifold corresponds to at least one stimulus. If each distinct stimulus also produces a unique population response, the mapping is one-to-one (injective). When the inverse mapping is also continuous, the neural response manifold is homeomorphic to the stimulus domain, meaning that they have the same topology, even though their Euclidean geometry can be different such as different curvatures and Euclidean distance. In addition, the tuning curves determine manifold geometry but not the reverse, so that the dimensionality and curvature of manifolds increase as tuning curves are narrowed. Finally, it has been shown that the geometry of the manifold determines both fine discrimination between neighboring coordinates and similarity between widely separated coordinates. In this study we use computational experiments to examine these connections for the contribution of color vision to characterizing fruits and vegetables. Usually, neural manifolds are estimated from neural state spaces that consist of the changes in population responses over time during a motor, perceptual or cognitive task. Instead of varying across time, the state space we examine will consist of changes in population responses as the input colors are changed.

The accurate decoding of single-color stimuli, achieved by a winner-take-all rule on the outputs of narrowly tuned IT cells [3,13], supported the notion that the color we see could be the preferred color of the IT neuron that fires the most. Narrowness of tuning can have different consequences for discrimination than identification depending on pair-wise correlations [29,30]. However, identification of fruits and vegetables that have color variations is inherently linked to discriminating between two color distributions, a problem that is quite different from discriminating between two colors, and more akin to separating at a later stage in a DNN, two manifolds that are intertwined or overlapping at the first stage [26].

## 2. METHODS

### 2.1 IT electrophysiology data

To model PIT responses, we used fMRI guided single-unit recordings done in alert monkey [9]. All cells in fMRI identified globs showed luminance-invariant color tuning. We used the responses of 228 spike-sorted cells to 20 out of the 45 test stimuli as these formed a set of equal luminance colors separated by roughly uniform color angles in CIELUV space [31] (Fig. 1A). To obtain continuous tuning curves for all colors, each cell’s mean responses were fit with an exponentiated-cosine tuning curve with the parameter *W* determining the permissible width of the deviation of the input color angle (*θ*) from the peak color (*θ_c_*), the parameter *m* the gradualness of the flanks, f_min_ the spontaneous response level and f_max_ the peak response [32] (Fig. 1B):

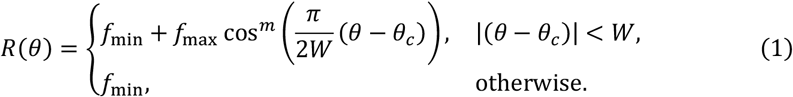

**Fig. 1.**
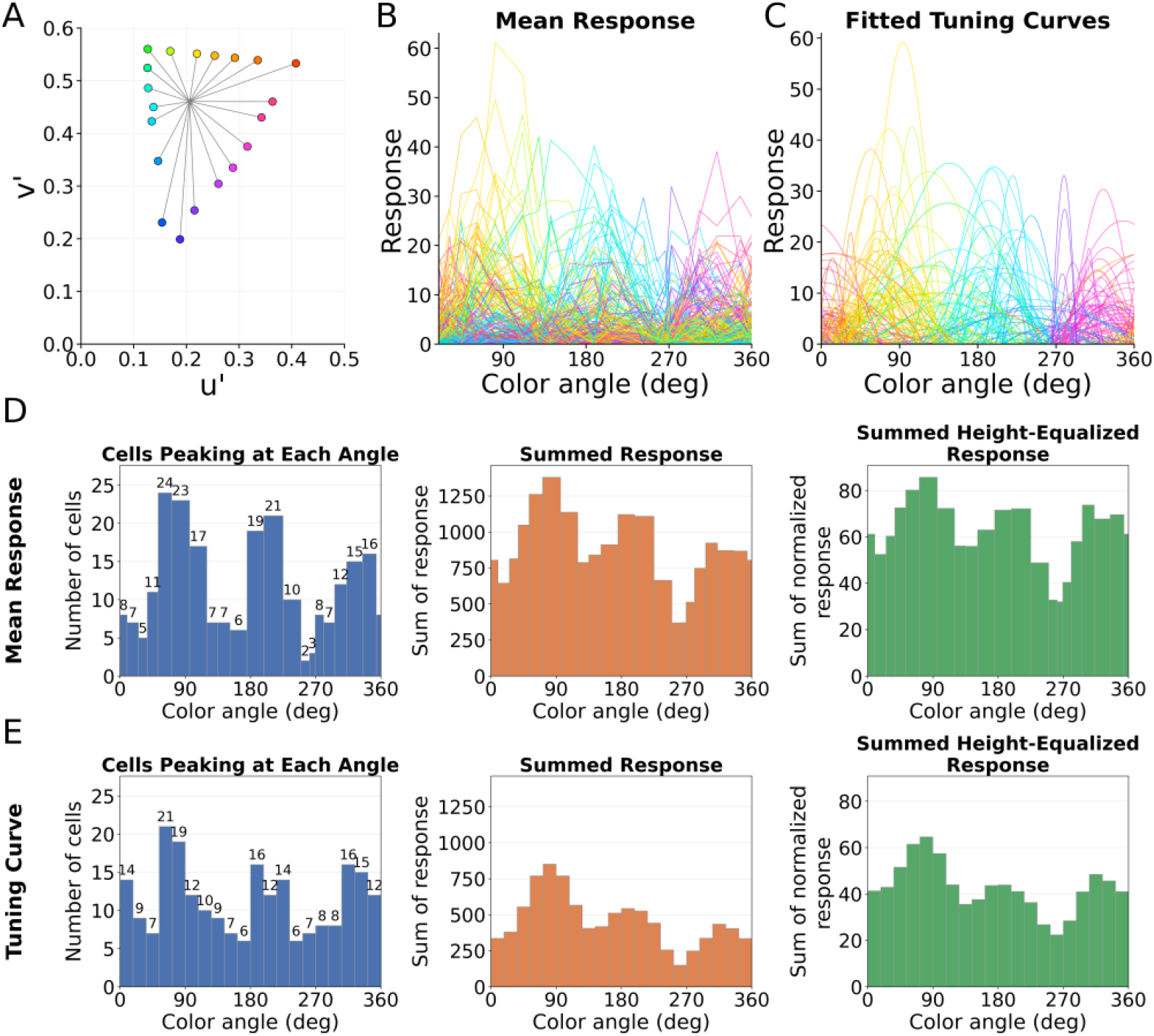
(A) 20 equal luminance color stimuli separated by approximately equal color angles in CIELUV space. (B) Mean and (C) fitted responses of 228 cells. The color of each curve corresponds roughly to the cell’s mean response peak. (D) Aggregated mean responses and (E) Responses of smooth tuning curves: Number of cells whose response peaks at each color angle, summed responses across the population at each color angle for the heights shown in panels B and C, and summed responses when heights are equated.

There were two differences between the calculations of the curves in Fig. 1C and the tuning curves previously fit to this data subset which was chosen as being approximately uniformly distributed in (u′, v′) color angles [31]. Almost equal spacing in color angle is the best that could be done post hoc with the data set, because it was not originally designed to estimate tuning curves. However, the distance from the mid-white point is not the same for each test color, so we equated the distances before curve fitting. We divided each response magnitude by the distance of its stimulus color from mid-white, relative to the shortest such distance. In essence, this creates a circle of colors with radius equal to the shortest distance of the original stimuli from the white point. In addition, we allowed the exponent of the cosine to be 1 or 2 to find the best fit for each tuning curve. The normalized mean responses and the fitted continuous tuning curves are fairly similar as shown in Figs. 1B and 1C. To examine the effect of the smoothing operation on population responses, Fig. 1E compared to Fig. 1D shows that smoothing the tuning curves smooths out the distributions of the number of cells peaking at each color angle, the summed response to each test color, and the summed height-equalized response to each test color but leaves the locations of peaks and valleys mostly unchanged. The stimulus set being restricted to a triangle in an iso-luminant plane, means that the tuning curves vary along just the color-angle dimension instead of being two or three dimensional and incorporating possible selectivity for saturation and brightness. Despite that limitation, we show what can be learned about IT by using computational comparisons to earlier neural stages.

### 2.2 Colors of 42 fruits and vegetables

The 42 fruits and vegetables we used together contain 4.5 million colored pixels (Fig. 2). The published dataset [20] contains measurements in CIE xyY, which we converted to L, M and S responses using the Smith-Pokorny cone fundamentals [33]. The dataset also has information in CIELUV space, which we converted to color angles to compute IT responses to the spectrum of each pixel.

**Fig. 2.**
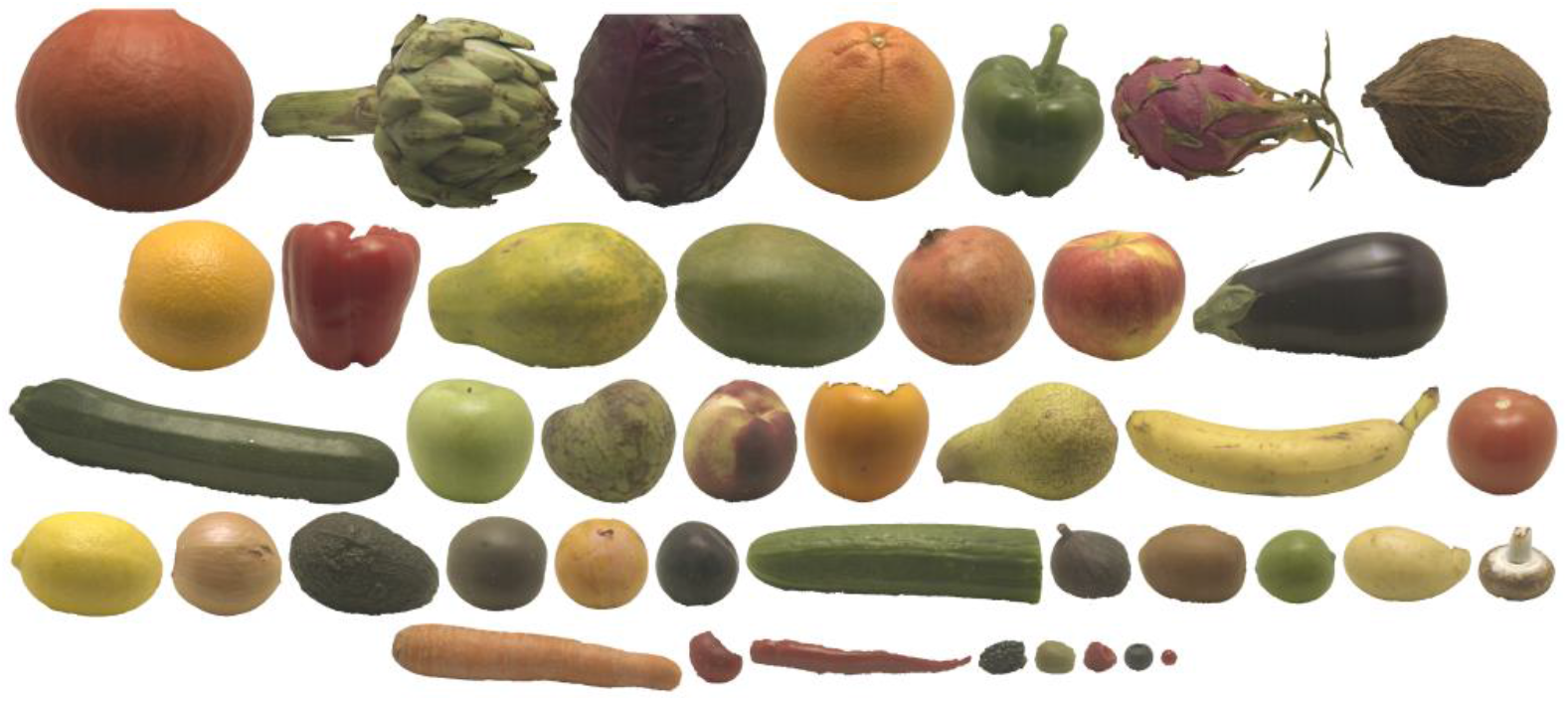
Hyperspectral images of 42 organic fruits and vegetables used to compute photoreceptor and IT responses. The full reflectance spectrum at every pixel was converted, under a fixed daylight illuminant (CIE 1931 xyY = [0.3324, 0.3435, 36.97 cd/m2]), into cone absorptions [20]. For display, we combined images of objects into a single montage that preserves their relative sizes

### 2.3 Classification of 42 fruits and vegetables using Fisher Discriminant Analysis

Because each downstream neuron receives input from a substantial population, it is natural to ask how much can be read out by a simple linear decoder. We used Fisher Discriminant Analysis (FDA) to determine the classification accuracy for different fruits and vegetables in the LMS and IT representations. For two sets of points, the FDA finds the best vector on which projecting the points maximizes the between-class variance and minimizes the within-class variance. The decision boundary is orthogonal to that vector. Assuming discrete classes of stimuli, FDA provides an upper bound on linear decoding capacity [34]

### 2.4 Adding Poisson variability

Neuronal recordings reveal that responses are noisy. A common way to characterize the random variability is to define each neuron as a Poisson device with the parameter for the mean and variance given by the tuning curve, which in turn is based on the average response to each test stimulus across repeated presentations [35]. *θ* is the color angle of the color stimulus. *r*_*i*_ is the number of spikes of neuron *i* in response to *θ* in a time interval. *R_i_*θ**:

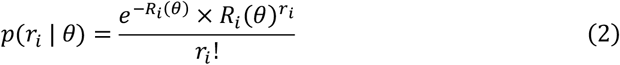

### 2.5 Visualizing the neural manifold using PCA and Isomap

Principal component analysis (PCA) is a linear dimensionality-reduction method: it projects the data onto orthogonal axes of maximal variance and therefore preserves Euclidean distances between points for the full dimensionality transform, but not for projection to reduced dimensions [36]. Isomap is a nonlinear method that instead preserves geodesic distances, measured along the surface of the data manifold [37]. PCA needs more dimensions to represent curved or folded manifolds while Isomap needs fewer dimensions and provides more interpretable manifold surface representations. We used both PCA and Isomap to understand IT population responses.

## 3. RESULTS

### 3.1 Manifolds of photoreceptor and IT responses to all color angles

To illustrate the notions of ambient neural space and reduced dimension manifolds, we first calculated the 3-D LMS state space consisting of L, M, S responses to 360 colors distributed uniformly around an equiluminant color circle in CIELUV space. Because cone absorption is a linear projection of the light spectrum, we used PCA to visualize the manifold which we expected to be 2-D. The first two principal components captured 100% of the variance confirming the expectation, and the colors formed the expected ellipsoid like their arrangement in the CIELUV chromaticity diagram (Fig. 3A).

**Fig. 3.**
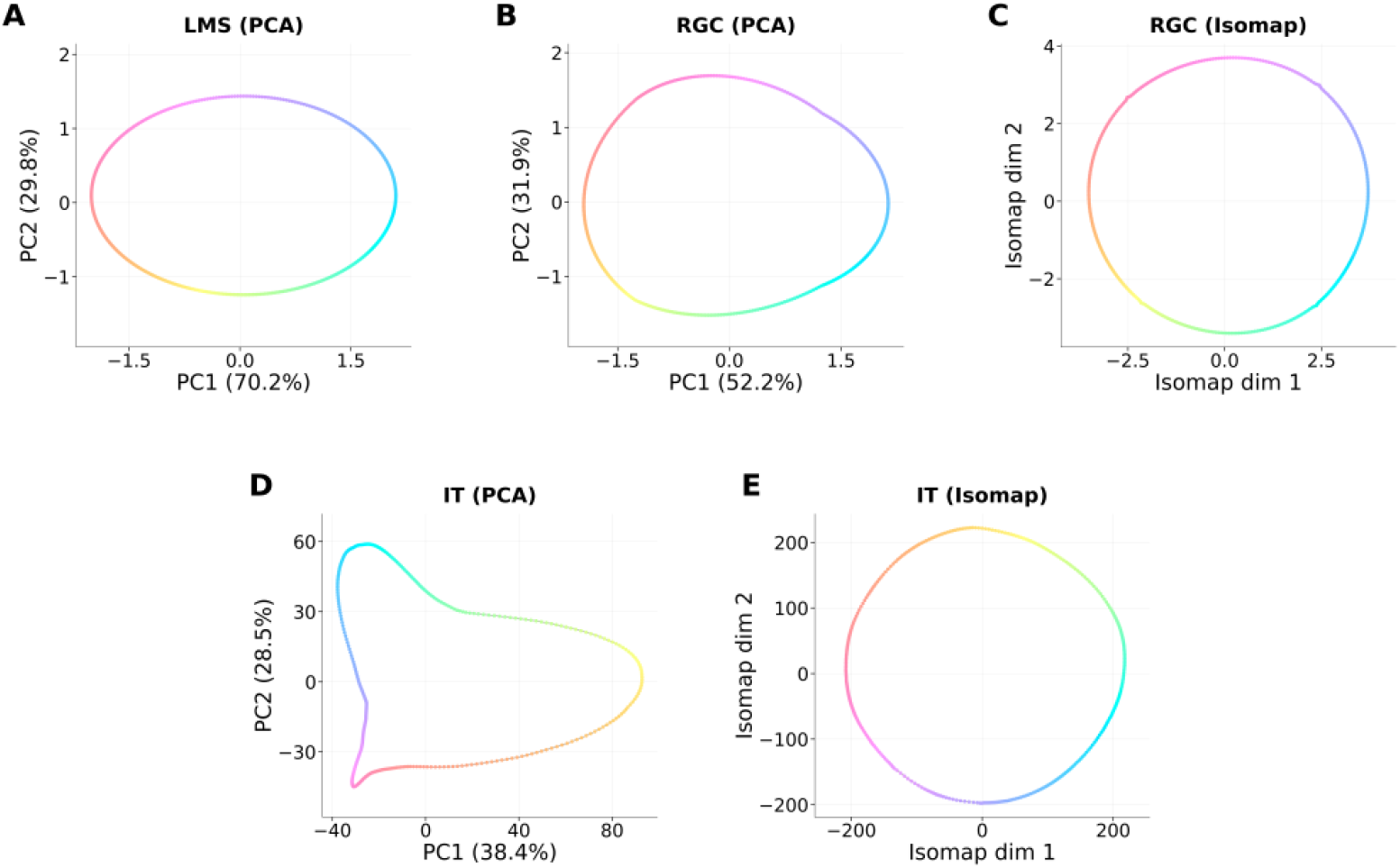
(A) PCA result of photoreceptors response on 360 stimuli colors. (B) PCA and (C) Isomap RGC 2-D manifolds for 360 colors. (D) PCA and (E) Isomap IT 2-D manifolds for 360 colors.

Between the cones and IT, however, color signals pass through several nonlinear processes. The first is the rectification into ON and OFF cells. To look at the effects of this operation we consider each polarity of the M and P retinal ganglion cells (RGCs) [11], forming four classes max[+(L − aM), 0.0], max[−(L − aM), 0.0], max[+(bS − (L + M)), 0.0], and max[−(bS − (L +M)), 0.0]. Using the coordinates of the mid-white point: (L_w_, M_w_, S_w_) = (2.0146, 1.0353, 0.0288), we calculated a = L_w_/M_w_ = 1.946 and b = (L_w_+M_w_)/S_w_ = 105.94, so that all four RGC classes are zero at the white point. We applied both PCA and Isomap to the RGC responses to the 360 colors. Across all hue angles, the first two principal components explain only 84% of the variance and the manifold is distorted into an egg shape (Fig. 3B). Isomap needed only two dimensions and yields a circular manifold (Fig. 3C).

We also applied both PCA and Isomap to the IT responses. In the PC1–PC2 plane, the hue trajectory in the IT manifold is distorted to a triangularish shape, stretched at yellowish, purplish, and bluish (Fig. 3D), reflecting the larger summed cell responses to these three color sectors due to the larger number of cells tuned to these sectors in this sample of neurons. If the heights of the tuning curves are equated, the manifold still shows a triangular shape but the corners get appreciably smoother and gradually curved. Across all hue angles, the first two principal components explain only 67% of the variance, and 8 components are needed to reach 95%. Isomap, in contrast, needs only two dimensions to represent the manifold. Consistent with the circular set of input colors, it orders all colors continuously along a closed loop (Fig. 3E). Previous applications of multidimensional scaling (MDS) to height-equated PIT responses to equiluminant colors gave results like the Isomap manifold [31,38]. Figure 3 demonstrates the usefulness of visualizing low-dimensional manifolds.

### 3.2 Photoreceptor responses to fruits and vegetables

To visualize the distribution of pixel color distributions across the objects, Fig. 4 shows the photoreceptor absorptions from the apple and orange pixels in three views of the 3-D LMS response space. Both sets of absorptions form narrow triangular fans with correlated L and M responses as is predictable from the overlap between the absorption spectra [39]. The two sets diverge at higher response magnitudes but have considerable overlap at lower magnitudes.

**Fig. 4.**
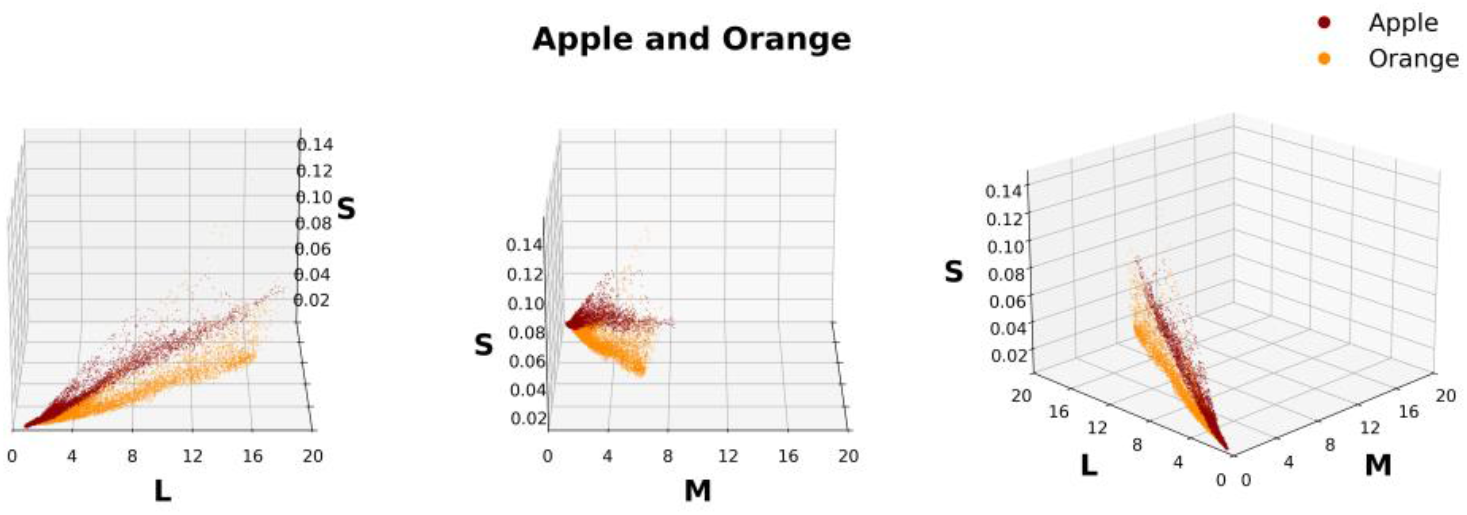
L, M, S response to each pixel in the apple and orange images.

To decorrelate the L and M responses, and to show the cone absorptions in a 2-D diagram, we have plotted the absorptions from each fruit and vegetable as L + M versus L − 2M, and the S absorption is indicated by the color of each point (Fig. 5). In these diagrams, the coordinates of the neutral illuminant were L + M = 3.05, L − 2M = −0.05 and S = 0.03. Points belonging to reddish fruits or vegetables lean towards the right side of the illuminant, while greenish fruits or vegetables lean toward the left. Although blue is less represented in fruits and vegetables, S-cone response to some of the pixels is visible in the panels. It is clear that points from some pairs of objects will have considerable overlap, making separation difficult with a linear separator if only cone responses or retinal ganglion cell responses [11] are compared. As suggested in earlier sections, IT cell responses can break up color space into discrete segments, thus the points from two objects could possibly be separated into regions of overlap and non-overlap, thus facilitating discrimination between objects with variegated colors solely on the basis of their color distributions. In the next section we test this possibility.

**Fig. 5.**
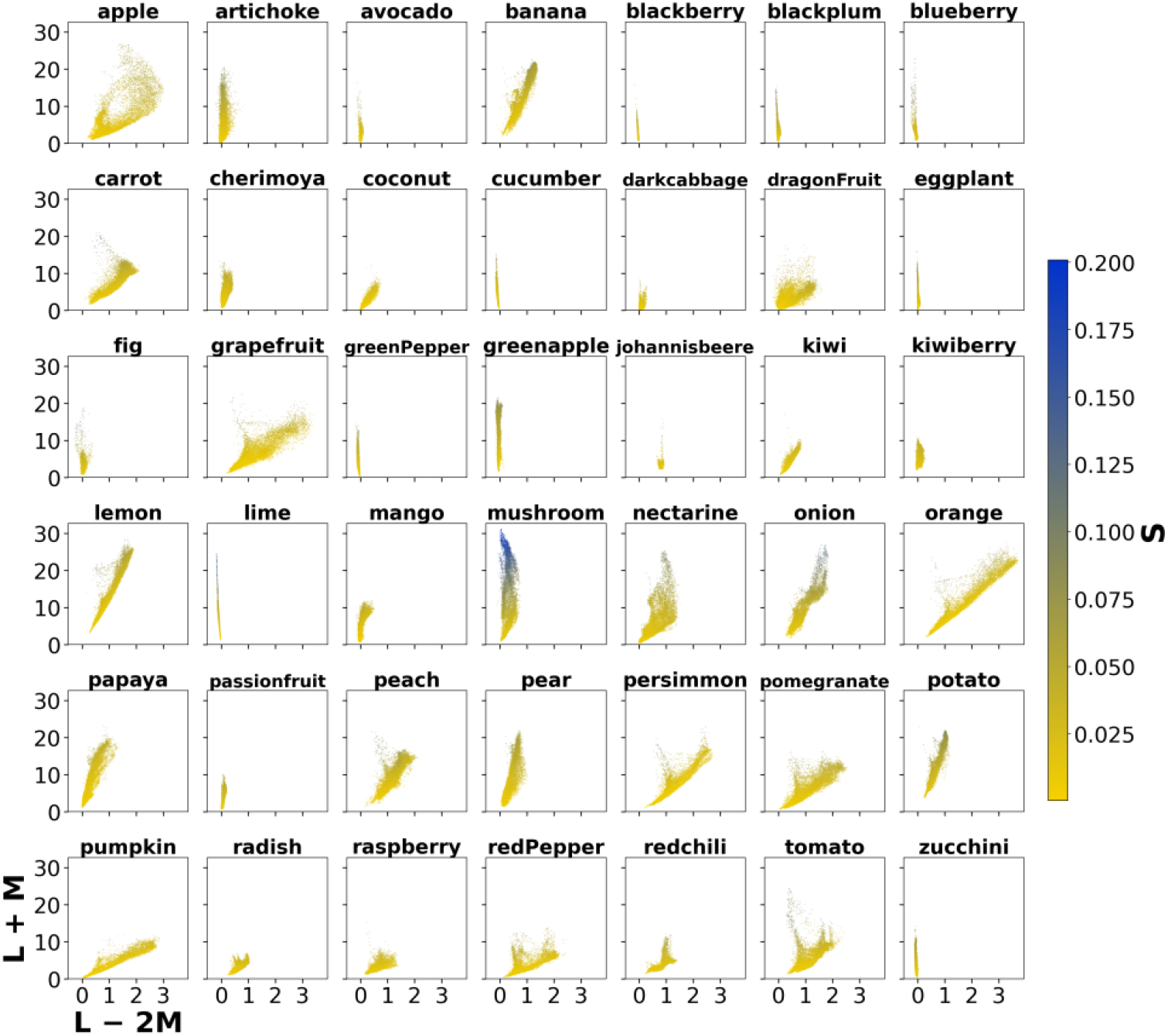
Luminance (L + M) versus L − 2M. The color bar shows the S cone response.

### 3.3 Fisher Discriminant Analysis of classification accuracy

Linear decoders are widely used in computational neuroscience because they approximate a single-layer readout by a downstream brain area, involving a weighted summation of inputs that indicates the information that is directly accessible by subsequent stages [40]. The explicit weights assigned to individual inputs make it easier to evaluate each unit’s contribution, although noise causes complications. The lower model complexity than non-linear models or DNNs reduces the risk of overfitting and provides a benchmark to compare different brain circuits or regions. Consequently, we have used linear Fisher Discriminant Analysis (FDA) to compare how well the populations of neural responses corresponding to different fruits and vegetables can be linearly separated at successive neural stages.

Figure 6 shows the results of doing pair-wise FDA on the LMS, RGC and IT neural population responses to fruits and vegetables. It is not clear what percentage of colors have to be separated for accurately classifying a pair, so we show the percentage of pairs for which 0– 100% of points can be separated. Without noise, all three stages are pretty successful and the improvement from LMS to RGC to IT is very slight. If there were no noise in the visual system, the narrow tuning curves in IT could facilitate identification of single colors but would not be needed to enhance separation of color distributions.

**Fig. 6.**
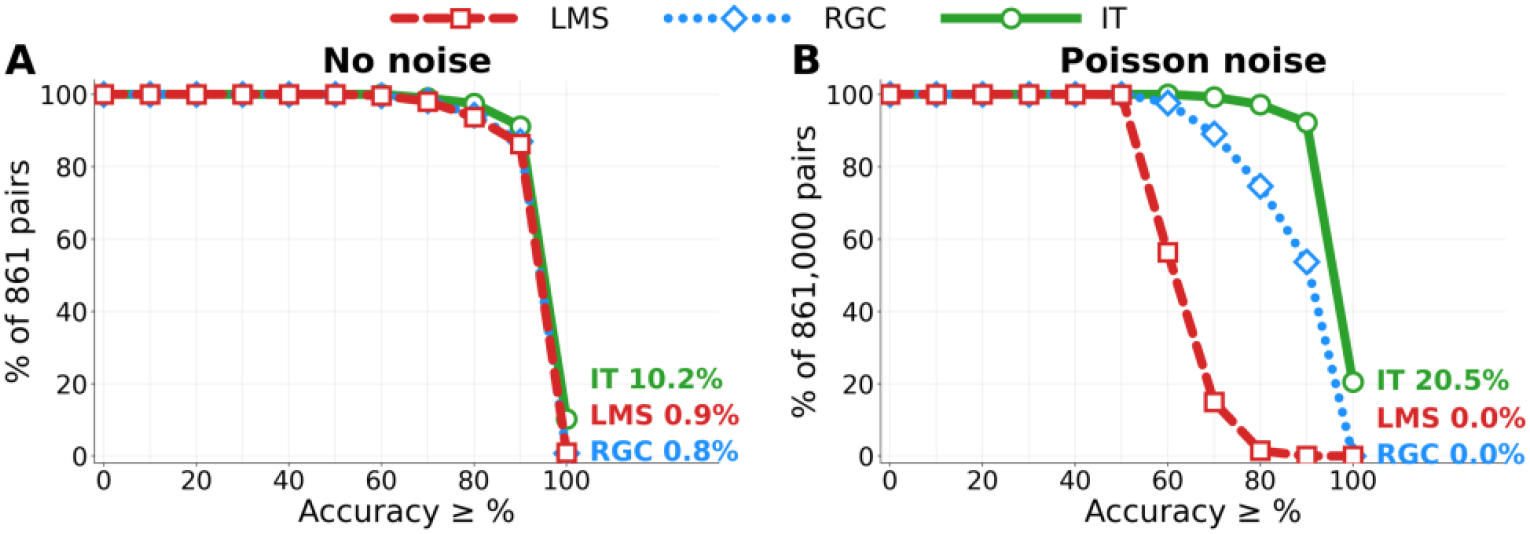
Result of Fisher Discriminant Analysis under (A) No noise and (B) Poisson noise conditions shown as the % of pairs for which at least 75%, 90% or 95% of points were separated.

A more realistic analysis includes random variability in neural responses. All stages of neural response exhibit some noise, due to multiple factors, but because noise does not keep increasing at successive stages, a model of successive stages would also have to incorporate noise reducing mechanisms such as integration [41–43]. To model noise in IT cells, we used Eq. (2), as we had done previously [3,13]. Noise in photoreceptors has multiple sources [44,45], and it has been modeled in multiple ways [46]. To simulate noise for photoreceptors that fairly matched the magnitude of IT noise, we took the maximum across all pixels of L, aM and bS responses, and set that value equal to the maximum number of spikes across all pixels of the 228 IT cells and then scaled the other photoreceptor responses proportionally. We set RGC noise similarly for the four classes of neurons. Since the variance of the Poisson distribution is equal to the mean, this method set roughly equal noise levels at the three stages. We then made 1000 simulations of fruit and vegetable manifolds with Poisson noise at each stage and performed pairwise FDA for each simulation. With Poisson noise, discrimination at the cone stage fell drastically (Fig. 6B), with only 5% of the pairs with 75% or more separated points, and no pairs with 90% or more separation. The situation improves considerably for RGC and even more at IT where 98%, 92% and 83% of pairs show separation of 75%, 90% and 95% of points. In the presence of realistic neuronal noise, the increased dimensionality of IT responses considerably enhances the ability to discriminate between natural distributions of colors.

In the Appendix, Tables A1 and A2 show the pairs of fruit and vegetables that improve the most in separation accuracy from LMS to IT, and the pairs that are hardest to separate at IT, both without considering noise. With simulated Poisson variability, IT separation is much better than LMS. While IT is better at separating objects with some overlapping colors like the apple and the carrot, it cannot separate objects like cucumber and green pepper that have similar color overall, even without noise. Figure A1 illustrates how some pairs get separation in the transformations from LMS to IT, and some do not, by using the overlap along the FDA axis (the orthogonal line is the decision boundary).

### 3.4 Neural manifolds of all fruits and vegetables for LMS, RGC and IT population responses

Although the FDA is done in 3-D space for LMS and 228-D space for IT, it is instructive to examine the 2-D PCA manifold for LMS, RGC and IT, and the 2-D Isomap manifold for RGC and IT for the combined response to all of the fruit and vegetable pixels (Fig. 7). The photoreceptor responses form a flat, two-dimensional triangular manifold, all three cone types contribute equally to the first PC, which explains 92.6% variance, and PC2 is similar to the S versus (L+M) axis where L and M have negative loadings, and S has positive loading. On the other hand, the response manifold in RGC is curved when the dimension is reduced by PCA. The main change is the separation of reds and greens as would be expected from the decorrelation of L and M responses [39,47], and that could explain a major component of the discrimination being better than LMS when noise is incorporated. For IT, the first 2 principal components explain 79.8% variance, and the first three explain 88.1% of the variance, reflecting the increased dimensionality of the population response. Inset in upper right corner shows the 3-D plot of the IT manifold. Each point in the plot is 228 IT neurons’ response to one pixel. Along PC1, reddish objects are on one side, and greenish objects are on the opposite side. Purple and yellow objects are on the opposite end of PC2. Since the 3-D PCA plot shows a curved 2-D plane embedding in the 3-D space, the space can be better visualized by nonlinear dimension reduction using Isomap. When it is flattened with Isomap, reddish objects fall at one end and greenish objects at the other, with yellowish and purplish objects likewise at opposite ends. In the LMS response manifold, the objects lie close together and overlap; in IT, due to higher dimensionality, they are more spread out. This greater separation predicts IT’s robustness to added Poisson neural variability.

**Fig. 7.**
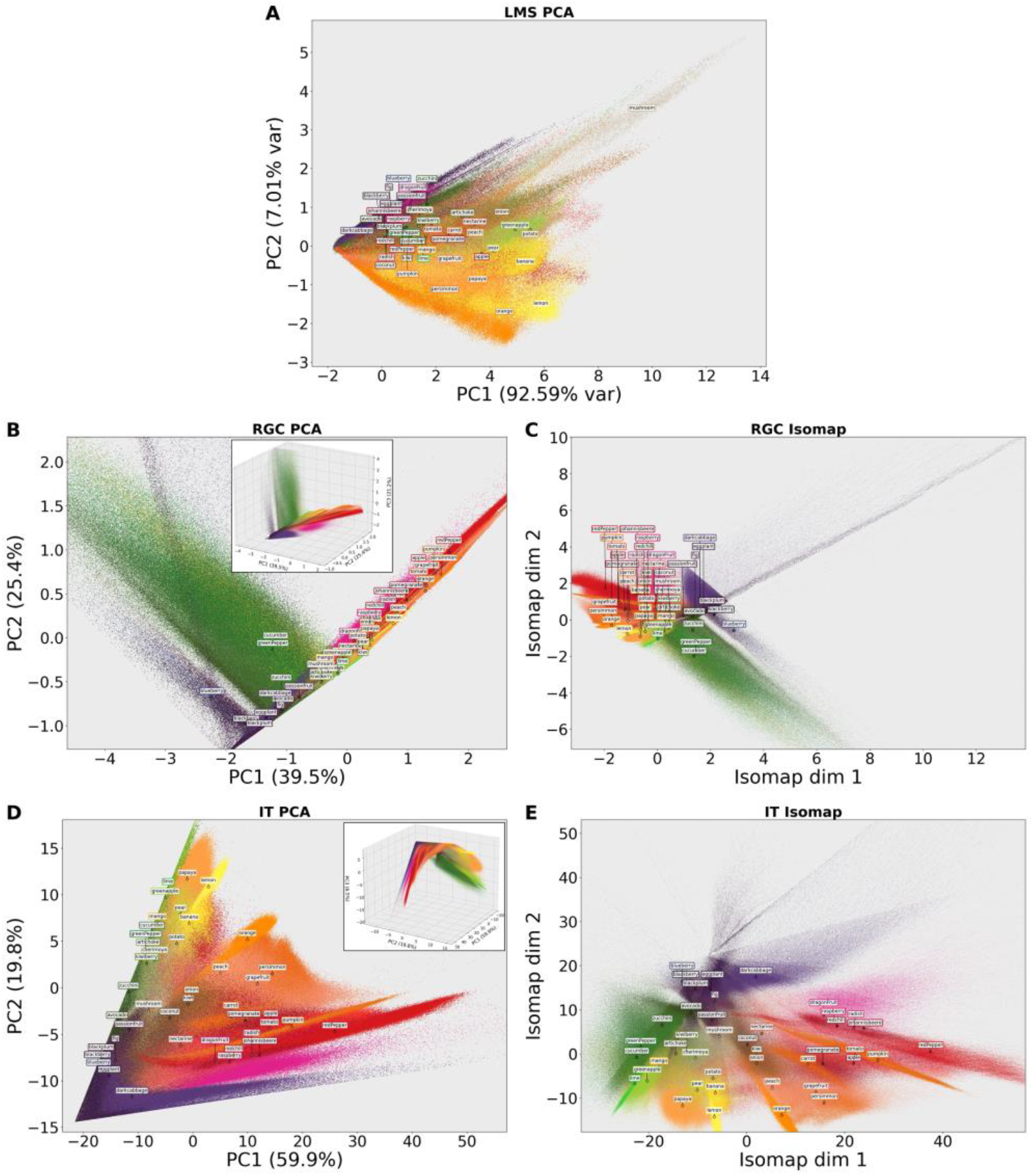
(A) PCA of LMS response manifold. (B) PCA result of RGC with 3-D plot in inset. (C) Isomap flattens the curved manifold, (D) PCA result of IT with 3-D plot in inset. (E) Isomap flattens the curved manifold.

## 4. DISCUSSION

Inferior temporal cortex encompasses a large area of cortex, serves many visual functions with some more or less specialized patches which are interconnected and possibly hierarchical [48]. Color is probably a part of many modalities that are represented in IT, but even for patches that seem devoted to color computations, much has to be learned about how color appearance is affected by categorization, spatio-temporal context, memory and language, etc. For example, besides pure color areas, Pennock *et al*. [49] found color-biased regions which respond strongly to food, beginning in V4 and diverging medially and laterally to the fusiform face area in both hemispheres, possibly using both color and form to represent food. Figuring out these functions of IT color responsive areas will require recording from populations of cells with specialized stimuli: 3-D tuning curves could be measured with responses as a function of phase for circular color excursions of different radii and orientations that modulate cone absorptions in sinusoidal fashion [11,50]. The effect of spatial context could be measured as color assimilation or induction by modulations being in the same or opposite phase as circular temporal modulations surrounding the receptive field with a uniform color and with a drifting spatial sinusoidal variation [51,52]. Whether a cell extracts transparency can be judged by estimating the extent to which color selectivity to the transparent layer is invariant to the color of the background [53], and if the degree of invariance corresponds to the subjective scale for transparency across motion, geometric and color cues [54]. Variations in the experimental conditions would also help in unraveling the processes with circuit models [55] complemented by analyses of manifolds using methods that have been worked out in other areas [56].

In this paper, we took on a simpler task, to understand one consequence of the narrowing and diversity of color responsive IT neurons. We went further than our earlier computations that used single colors as stimuli, to looking at the discrimination between the distributions of colors that define natural fruits and vegetables. We showed that with realistic neuronal response variability, the population of IT neurons is better at such discriminations than would be expected from just cone outputs. We are aware that this is not quite a natural task, because we are throwing out information about the spatial gradient of colors, and the 3-D shape of the object, but that simulations show that it is a difficult task to accomplish with just noisy cone responses, but is facilitated by the high-dimensionality of IT. That IT cannot separate every pair successfully could be expected, given that there are pairs of these color distributions (Fig. 8) that humans cannot discriminate either.

**Fig. 8.**
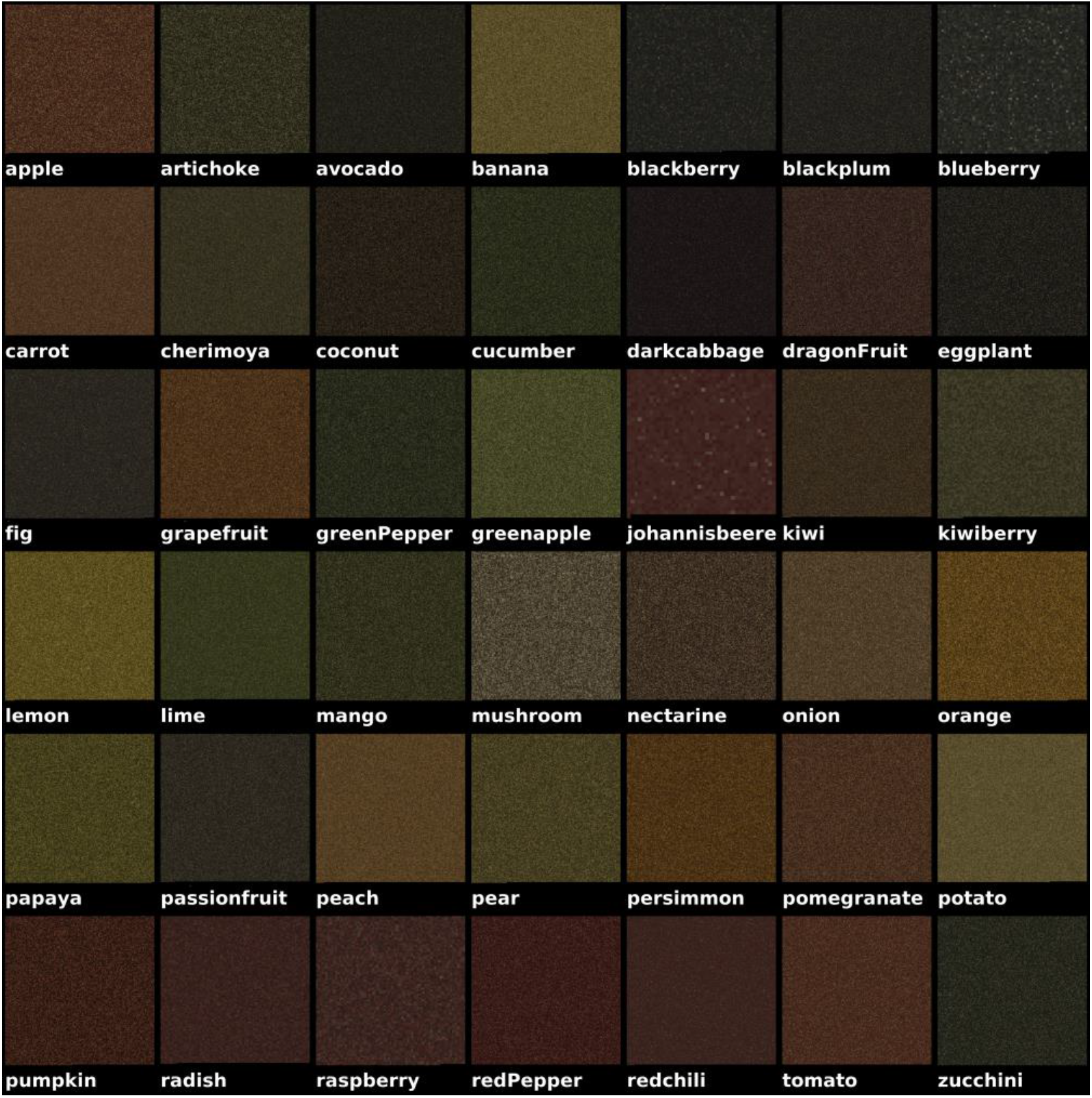
Each square shows a spatially randomized arrangement of all the color pixels of the labelled vegetable or fruit from the images in Fig. 2.

## Disclosures

The authors declare no conflicts of interest.

### Data availability

The physiology data were published in [9]. The hyperspectral dataset of fruits and vegetables was published in [20].

## APPENDIX

**Table A1.** Fruit and vegetable pairs whose classification accuracy improves the most from the photoreceptors (L, M, S) to IT. Accuracy is the percentage of pixels correctly separated by pairwise Fisher Discriminant Analysis without noise; the 20 of 861 pairs that improve by more than 10 percentage points are listed.

| **Fruit pair** | **LMS (%)** | **IT (%)** | **IT − LMS (%)** |
|---|---|---|---|
| apple – carrot | 64.0 | 91.2 | +27.1 |
| coconut – dragonFruit | 70.4 | 94.4 | +23.9 |
| apple – peach | 68.9 | 90.3 | +21.5 |
| dragonFruit – kiwi | 74.6 | 93.4 | +18.9 |
| apple – redPepper | 69.7 | 88.3 | +18.6 |
| apple – pomegranate | 60.5 | 78.6 | +18.2 |
| nectarine – onion | 72.1 | 87.7 | +15.7 |
| avocado – eggplant | 66.7 | 81.7 | +14.9 |
| coconut – pumpkin | 86.8 | 100.5 | +13.7 |
| artichoke – coconut | 83.8 | 97.0 | +13.2 |
| dragonFruit – pumpkin | 85.4 | 98.5 | +13.1 |
| nectarine – redchili | 77.3 | 89.6 | +12.2 |
| avocado – nectarine | 85.6 | 97.0 | +11.5 |
| apple – grapefruit | 78.5 | 89.9 | +11.4 |
| darkcabbage – dragonFruit | 88.7 | 99.8 | +11.1 |
| greenapple – lime | 79.6 | 90.6 | +11.0 |
| kiwi – nectarine | 76.1 | 87.1 | +11.0 |
| coconut – eggplant | 85.1 | 95.9 | +10.9 |
| apple – pumpkin | 78.9 | 89.5 | +10.5 |
| greenPepper – nectarine | 90.4 | 100.9 | +10.5 |

**Table A2.** Fruit and vegetable pairs that are hardest to separate at IT: the 20 of 861 pairs with the lowest percentage of pixels correctly separated by pairwise Fisher Discriminant Analysis of the IT responses without noise.

| **Fruit pair** | **IT (%)** |
|---|---|
| kiwi – onion | 63.7 |
| cucumber – greenPepper | 63.9 |
| johannisbeere – radish | 64.2 |
| radish – raspberry | 65.2 |
| apple – tomato | 65.5 |
| coconut – kiwi | 67.7 |
| johannisbeere – raspberry | 68.8 |
| blackberry – blackplum | 69.9 |
| radish – redchili | 70.5 |
| grapefruit – persimmon | 71.0 |
| coconut – onion | 72.2 |
| blackberry – blueberry | 72.8 |
| carrot – pomegranate | 72.9 |
| artichoke – kiwiberry | 73.4 |
| artichoke – cherimoya | 73.8 |
| artichoke – mango | 76.1 |
| dragonFruit – nectarine | 76.5 |
| cherimoya – kiwiberry | 78.2 |
| raspberry – redchili | 78.4 |
| apple – pomegranate | 78.6 |

**Fig. A1.**
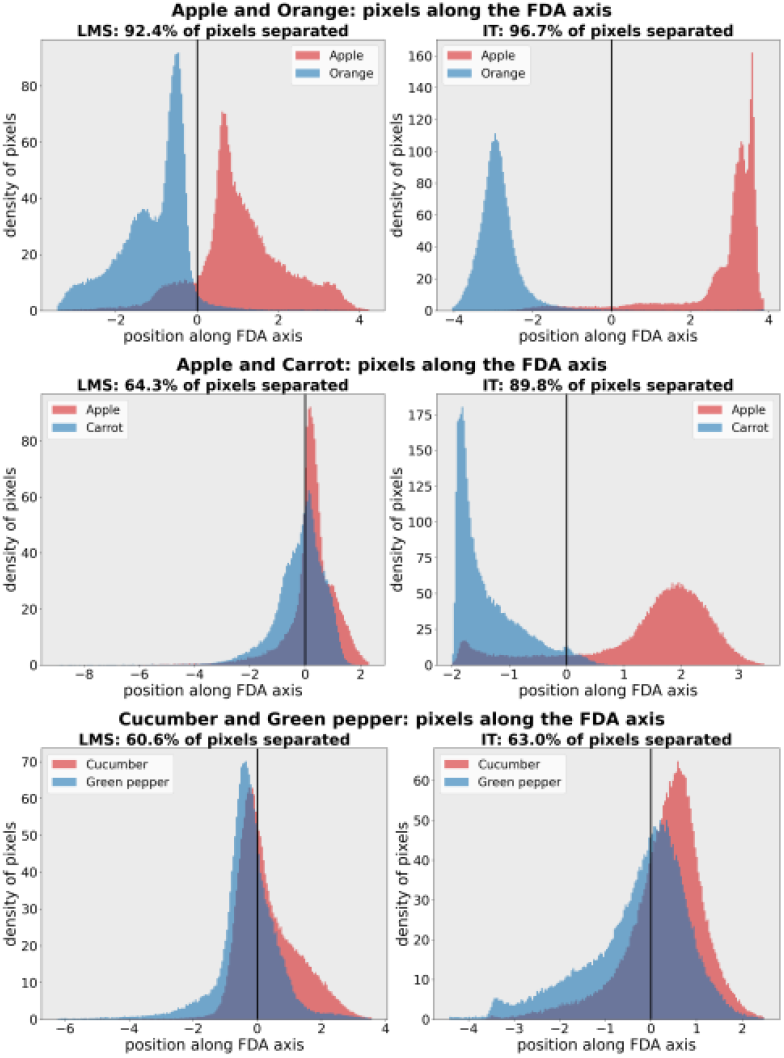
Example of separable, hard to separate, best separation improvement pair from the photoreceptors to IT. The plot shows linear boundary that separates pairs of objects. It projected L, M, S and IT response along the FDA vector. While apple and orange can be separated at the photoreceptors, the separation becomes cleaner at IT. The improvement of separation is even larger for apple and carrot. Cucumber and green pepper are hard to separate both in L, M, S and IT.

